# Tractography Reproducibility Challenge with Empirical Data (TraCED): The 2017 ISMRM Diffusion Study Group Challenge

**DOI:** 10.1101/484543

**Authors:** Vishwesh Nath, Kurt G. Schilling, Prasanna Parvathaneni, Allison E. Hainline, Yuankai Huo, Justin A. Blaber, Matt Rowe, Paulo Rodrigues, Vesna Prchkovska, Dogu Baran Aydogan, Wei Sun, Yonggang Shi, William A. Parker, Abdol Aziz Ould Ismail, Ragini Verma, Ryan P. Cabeen, Arthur W. Toga, Allen T. Newton, Jakob Wasserthal, Peter Neher, Klaus Maier-Hein, Giovanni Savini, Fulvia Palesi, Enrico Kaden, Ye Wu, Jianzhong He, Yuanjing Feng, Muhamed Barakovic, David Romascano, Jonathan Rafael-Patino, Matteo Frigo, Gabriel Girard, Alessandro Daducci, Jean-Philippe Thiran, Michael Paquette, Francois Rheault, Jasmeen Sidhu, Catherine Lebel, Alexander Leemans, Maxime Descoteaux, Tim B. Dyrby, Hakmook Kang, Bennett A. Landman

## Abstract

**Purpose:** Fiber tracking with diffusion weighted magnetic resonance imaging has become an essential tool for estimating in vivo brain white matter architecture. Fiber tracking results are sensitive to the choice of processing method and tracking criteria. Phantom studies provide concrete quantitative comparisons of methods relative to absolute ground truths, yet do not capture variabilities because of in vivo physiological factors.

**Methods:** To date, a large-scale reproducibility analysis has not been performed for the assessment of the newest generation of tractography algorithms with in vivo data. Reproducibility does not assess the validity of a brain connection however it is still of critical importance because it describes the variability for an algorithm in group studies. The ISMRM 2017 TraCED challenge was created to fulfill the gap. The TraCED dataset consists of a single healthy volunteer scanned on two different scanners of the same manufacturer. The multi-shell acquisition included b-values of 1000, 2000 and 3000 s/mm^2^ with 20, 45 and 64 diffusion gradient directions per shell, respectively.

**Results:** Nine international groups submitted 46 tractography algorithm entries. The top five submissions had high ICC > 0.88. Reproducibility is high within these top 5 submissions when assessed across sessions or across scanners. However, it can be directly attributed to containment of smaller volume tracts in larger volume tracts. This holds true for the top five submissions where they are contained in a specific order. While most algorithms are contained in an ordering there are some outliers.

**Conclusion:** The different methods clearly result in fundamentally different tract structures at the more conservative specificity choices (i.e., volumetrically smaller tractograms). The data and challenge infrastructure remain available for continued analysis and provide a platform for comparison.

## 1. INTRODUCTION

Diffusion weighted magnetic resonance imaging (DW-MRI) is a technique which allows for non-invasive mapping of the human brain’s micro-architecture at milli-metric resolution. Using voxel-wise fiber orientation reconstruction methods, tractography can provide quantitative and qualitative information for studying structural brain connectivity and continuity of neural pathways of the nervous system in vivo. There have been many algorithms, global, iterative, deterministic and probabilistic, that reconstruct streamlines using fiber reconstruction methods. Tractography was conceived [2] using one of the first fiber reconstruction method, diffusion tensor imaging (DTI) [1]. However, DTI has a well-known limitation: it cannot resolve complex fiber configurations [3]. With the advancement in acquisitions protocols allowing for better resolution and higher number of gradient values new methods for reconstruction of local fiber have been developed. These methods are commonly referred to as high angular resolution diffusion imaging (HARDI), e.g., q-ball, constrained spherical deconvolution (CSD), persistent angular structure (PAS) [4-6]. HARDI methods enable characterization of more than a single fiber direction per voxel, but have been often shown to be limited when more than two fiber populations exist per voxel [7, 8]. While there is definite gain in sensitivity when using HARDI methods, there remain critical questions of their reproducibility [9].

There have been many validation efforts that aim to assess the anatomical accuracy of tractography. Early studies investigated how well tractography followed large white matter trajectories through qualitative comparisons with dissected human samples [10], or previous primate histological tracings [11]. Later works on the macaque [12] or porcine [13] brains highlighted limitations and common errors in tractography. Recently, the sensitivity and specificity of tractography in detecting connections has been systematically explored against tracers in the monkey [14-16], porcine [17], or mouse [18] brains. The main conclusions drawn from these are (1) that algorithms always show a tradeoff in sensitivity and specificity (i.e. those that find the most true connections have the most false connections) (2) short-range connections are more reliably detected than long-range, (3) connectivity predictions do better than chance and thus have useful predictive power, and (4) tractography performs better when assessing connectivity between relatively large-scale regions rather than identifying fine details or connectivity.

Despite the wide range of validation studies, there have been few reproducibility studies of tractography [19-21]. Rather than ask how right (or wrong) tractography is, we ask how stable are the outputs of these techniques? Because tractography is an essential part of track segmentation, network analysis, and microstructural imaging, it is important that reproducibility is high, otherwise power is lost in group analyses or in longitudinal comparisons. In this study, given a standard, clinically realistic, diffusion protocol, we aim to assess how reproducible tractography results are between repeats, between scanners, and between algorithms.

Publicly organized challenges provide unique opportunities for research communities to fairly compare algorithms in an unbiased format, resulting in quantitative measures of the reliability and limitation of competing approaches, as well as potential strategies for improving consistency. In the diffusion MRI community, challenges have focused on recovering intra-voxel fiber geometries using synthetic data [22] and physical phantoms [19, 23]. Similarly, diffusion tractography challenges [20] have provided insights into the effects of different acquisition settings, voxel-wise reconstruction techniques, and tracking parameters on tract validity by comparing results to ground truth physical phantom fiber configurations [19, 21]. Recently, more clinically relevant evaluations have been put forth. For example, a recent MICCAI challenge benchmarked DTI tractography of the pyramidal tract in neurosurgical cases presenting with tumors in the motor cortex [24]. Towards this direction, the current challenge utilized a large-scale single subject reproducibility dataset, acquired in clinically feasible scan times. This challenge was intended to study reproducibility to describe the limitations for capturing physiological and imaging considerations prevalent in human data and evaluate the newest generation of tractography algorithms.

This paper is organized as follows. First, we present the analysis structure of this challenge to characterize which tracts are the most reproducible. Second, we characterize the variance across the tractography methods by design features and compare the potential containment of tracts on a per algorithm basis.

## 2. METHODS

### 2.1 DW-MRI Data Acquisition

The data were acquired with a multi-shell HARDI sequence on single healthy human subject. The two scanners were both Phillips, Achieva, 3T, Best, Netherlands. These are referred to as scanner ‘A’ and ‘B’. The three shells that were acquired: b=1000 s/mm^2^, 2000s/mm^2^ and 3000s/mm^2^ with 20, 48 and 64 gradient directions respectively (uniformly distributed over a hemi-sphere and independently per shell, this was done in consideration of scanner hardware.). The other parameters were kept consistent for all shells. They are as follows: Delta=∼48ms, delta=∼37ms, partial fourier=0.77, TE = 99 ms, TR ∼= 2920 ms and voxel resolution=2.5mm isotropic. A total of 15 non-weighted diffusion volumes ‘b0’ images interspersed as 5 per shell were acquired. Additionally, for scanner A & B, 5 reverse phase-encoded b0 images and 3 diffusion weighted directions were acquired to aid in distortion correction. The additional 3 diffusion-weighted direction volumes were acquired for ease of acquisition from the scanner. They do not contribute to the pre-processing of the data in any way.

Additionally, a T1-weighted reference image (MPRAGE) was acquired for each session per scanner (4 volumes total). A single volume of T1 was used which was registered to the first session of scanner A where the session had already been registered to the MNI template. This was done using a 6 degree of freedom rigid body registration.

For the initial data release, a technical issue resulted in 5 *non-*reverse phase-encoded b0 images for scanner A. Note that at the end of the challenge, the scanner ‘A’ data were completely re-acquired for both sessions with 5 reverse phase-encoded b0 images and 3 diffusion weighted directions. These data were released as supplementary material, but not included in the presented challenge data. Following the protocol for tractography in [25], we delineated six tracts cingulum (CNG) Left/Right (L/R), inferior longitudinal fasciculus (ILF) (L/R), inferior fronto-occipital (IFO) (L/R). The mean intra-class correlation (ICC) inter-scanner values for the original challenge data and the updated challenge data were 0.86 and 0.89, respectively. The mean difference between methods was 0.15 in terms of ICC. As expected, the inclusion of full reverse phase encoding for Scanner ‘A’ introduced a small increased in consistency relative to much larger differences between methods.

DW-MRI Data Pre-processing as illustrated in Fig 1, the 5 repeated acquisitions from each of the four sessions (two repeated on scanner A and B) were concatenated and corrected with FSL’s eddy and topup [25-27]. Intensity normalization was performed by dividing each diffusion weighted scan by the mean of all non-weighted diffusion volume (B0) per session. The average B0 from scanner A of the first sessions was rigidly (six degrees of freedom) registered [28] to a 2.5 mm T2 MNI template (this was done to ensure resampling from registration was done on both datasets). Next, the average B0 from the scanner A second session was rigidly registered to the average B0 of the registered scanner A first session B0 which had already been registered to the MNI space. Successively, the sessions from scanner B were registered to the sessions of scanner A. The b-vectors were rotated to account for the registration of the DW-MRI data [29].

**Figure 1:**
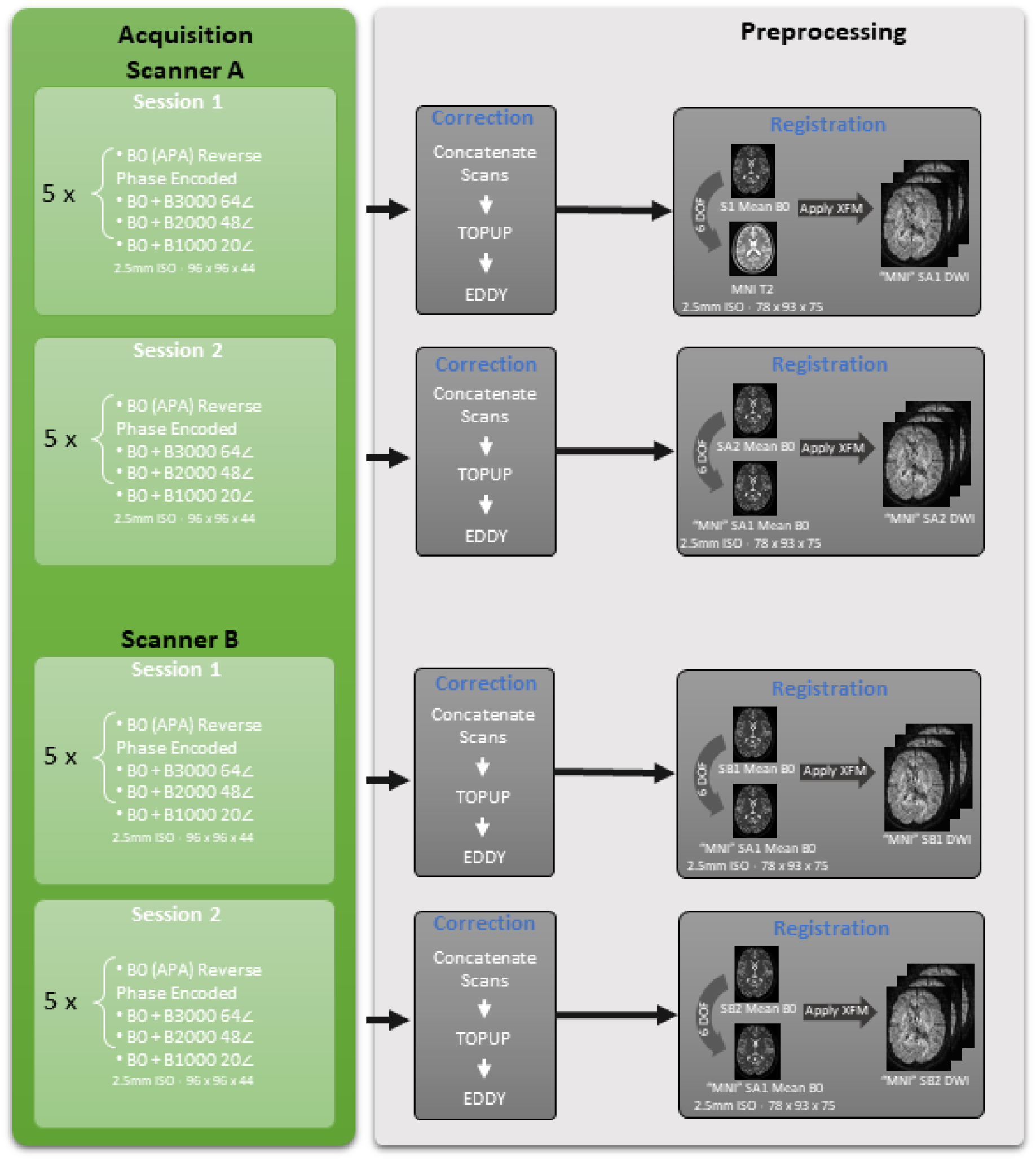
The acquisition per session included five repeats of a single b0 and successively at b-values of 3000, 2000 and 1000 s/mm^2^ using 64, 48 and 20 gradient directions respectively. Each session was individually corrected using topup, eddy and then normalized. All the sessions were registered using flirt to the first session of scanner A.

The T1 weighted MPRAGE was rigidly registered to the average registered b0 from the first session of scanner A. This transformation was applied to the T1 maintaining 1 mm isotropic resolution, thus providing a high-resolution segmentation that may be converted into diffusion space by performing a simple down-sampling. Multi-atlas segmentation with non-local spatial STAPLE fusion was used for the segmentation of the T1 volume to 133 different ROI’s [30, 31]. Finally, Multi-atlas CRUISE (MaCRUISE) was used to identify cortical surfaces [32]. These were provided for ease of algorithm implementations.

An informed consent under the Vanderbilt University (VU) Institutional Review Board (IRB) was obtained to conduct this study.

### 2.2 Challenge Rules and Metrics

For each of the 20 HARDI datasets (5 repetitions x 2 sessions x 2 scanners), participants were asked to submit a tractogram (i.e., “fiber probability membership function”) for each well-modeled fiber structures (uncinate (UNC) [L/R], fornix (FNX) [L/R], genu of the corpus callosum, cingulum (CNG) [L/R], corticospinal tract (CST) [L/R], splenium of the corpus callosum, inferior longitudinal fasciculus (ILF) [L/R], superior longitudinal fasciculus (SLF) [L/R], and inferior fronto-occipital (IFO) [L/R](1)). Each tractogram is a NIFTI volume at the field of view and resolution of the T1-weighted reference space where the floating-point value (32-bit single precision) of each voxel is in [0, 1] and indicates the probability of the voxel belonging to the specified fiber tract. Thus, participants submitted a total of 320 (5 x 2 x 2 x 16) NIFTI volumes using the acquisition of both the scanners. Assessment of fiber fractions was supported (i.e., the sum across all tracts is <=1 with the remainder as background). However, strict probabilities where each voxel may have a high probability of 2 or more fibers with a sum greater than 1 were permitted as well.

Tractograms within a submission were compared based on reproducibility of the tracts (intra-class correlation coefficient (ICC) statistics for continuous values and Dice similarity scores based on maximum probability assignment at 0.5). Intra-session, inter-session, same scanner, and inter-scanner scanner metrics have been reported for quantitative interpretation. The ICC and dice value of unique number of combinations of pairs of repeats were used as data points for violin plots depicting results of intra-session, inter-session and inter-scanner. The unique combinations of repetitions were 40, 50 and 100 respectively for the three levels of reproducibility.

### 2.3 Containment Analysis

A key question is whether the differences in tractography are driven by different considerations of the volume of the track, i.e., the larger the volume is, the more likely the track may include the underlying true track. For example, it is plausible that a set of tractography methods could see the same underlying probabilistic connection pattern and choose to threshold it based on different preferences for the volume of tracks. If the preference was driving the tractography differences, then tractograms would essentially be able to be nested from smallest to largest. To examine this hypothesis, we define the property containment index (CI) for two tracts where

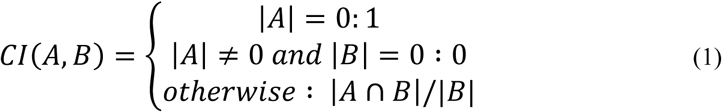

For the purposes of this discussion, we define the tractogram set to be the binary volume resulting at a 0.05 threshold of the mean of all results submitted for each algorithm. A visual understanding of containment index can be observed in Fig 3.

Then, an optimal ordering (“nesting”) of tractogram entries can be computed by maximizing the containment energy (CE, i.e., sum of CI for all tracts versus the tracts earlier than the one under consideration):

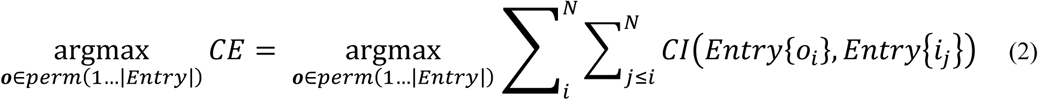

Where perm denotes the permutation operator and Entry is a list of all entered tractograms. Conceptually, this procedure finds the ideal order to stack the tractograms inside each other where the first tract is “most inside” the subsequent ones and the last tract is “most outside” all others. We define <CI> as the average containment index of all nesting for the ordered entries that are smaller than or equal to an entry provides a quantitative way to examine “nesting” (note, this approach includes the self-containment index so that the first entry has a CI of 1). Then, we can see how the nesting holds up from the inner (#‘1’) to the outer (#‘46’) entry.

## 3. RESULTS

Table 1 presents a more detailed technical contribution of each of the works:

- Team 1, Team 5, Team 6, Team 8 and Team 9 used all three shells of b-values provided in the dataset. Team 2 used all shells with data from an additional 30 subjects from the Human Connectome Project. Team 3 used shells of b-values 1000 and 2000 s/mm^2^. Team 7 and Team 4 only used the shell of b-value 3000 s/mm^2^.
- Additional pre-processing has been used by four teams. Team 4: Data was up-sampled to 1mm isotropic resolution. Team 6 used image de-noising techniques and up-sampled the data to 1.25mm. Team 5 and Team 9 used different styles of segmentation of the data presented for analysis.
- In terms of the fiber detection model, Team 6 and Team 3 used variants of tensor models while the others have used different variants of constrained spherical deconvolution. Notably Team 8 used a compartment analysis model using spherical harmonics.
- Considering the tractography parameters - the range of step sizes that have been used lie between 0.2-1.25mm. Threshold angle lies in the range of 20-40 degrees.
- Single fiber assumptions were considered with the condition of FA > 0.7 by teams Team 1 and Team 4. A notable observation here is that a general assumption was made by Team 6 to reject voxels which were less than 0.15 FA.
- Team 2, Team 6 and Team 8 post-processed the tractography results for removal of spurious fibers by defining different and specific constraints.
- Of note, Team 2 treated the tractography problem as a segmentation problem and developed a U-net which was trained on the HCP data. While Team 9 used a multi-atlas approach to tractography. The other teams used the general approach of probabilistic or deterministic tractography.

**Table 1:**
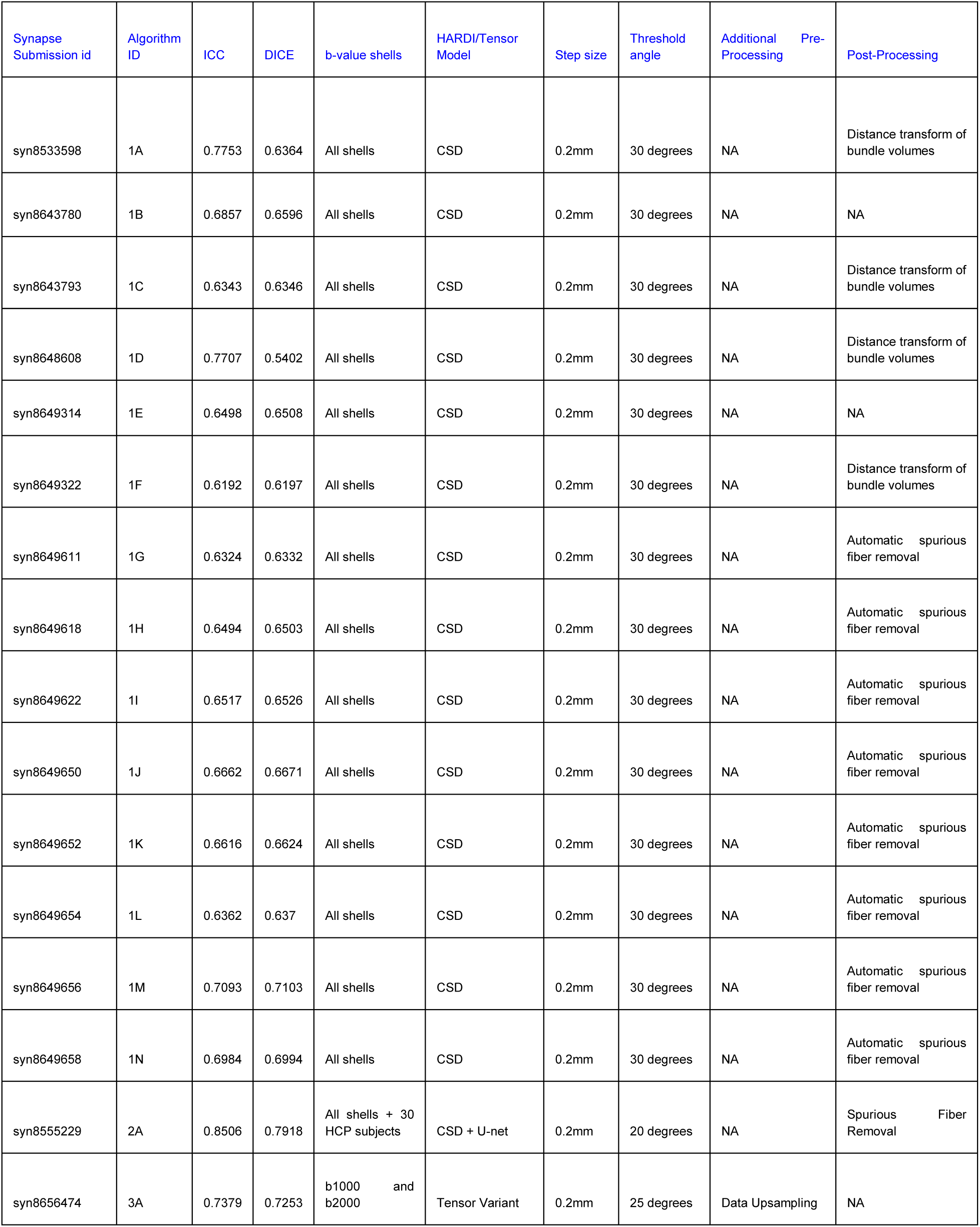
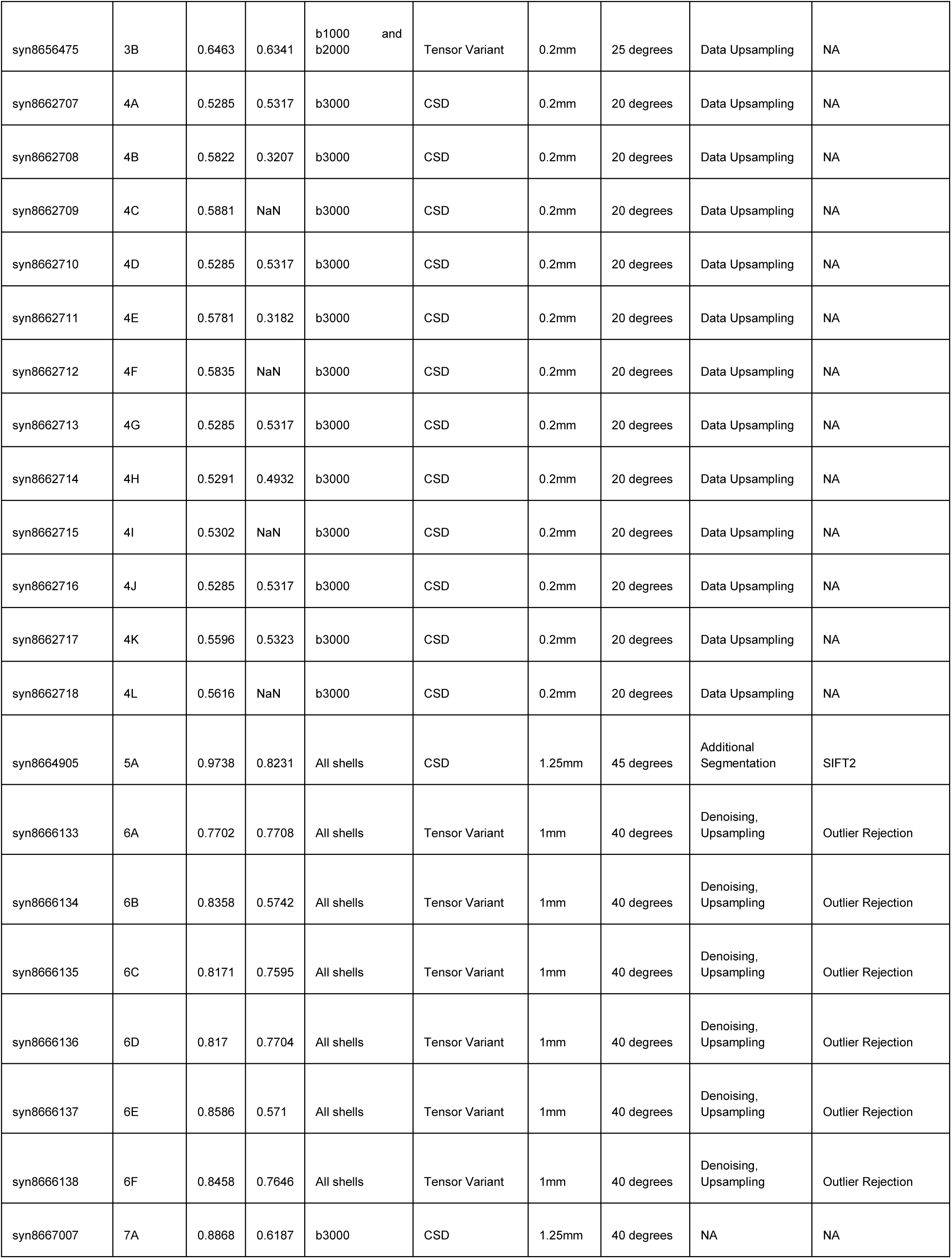
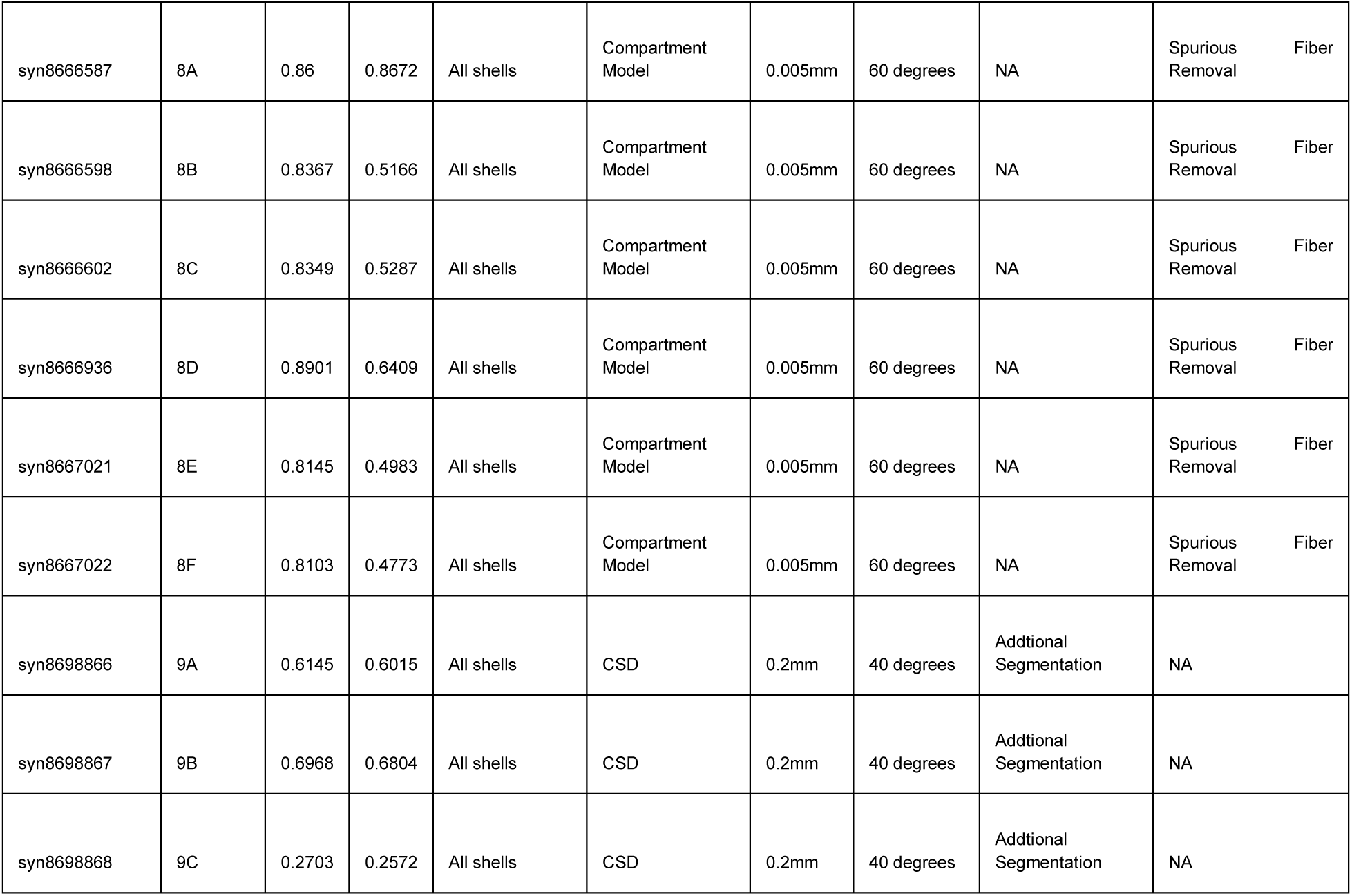
The table presents all the hyper-parameters of the different algorithms that were submitted and an overall evaluation of the algorithm in terms of ICC and Dice.

An overlay of all 46 submissions, for all estimated fiber pathways can be observed (Fig 2 Column 1 & 3). Only the left side has been shown as the right side is a similar observation. There are vast differences that can be noticed in the estimated pathways. The volume of the brain occupied by each tract from different submissions varied dramatically. When all 46 submissions are overlaid, tracts occupy 14-53% of the brain volumetrically (average – 34%). Specifically, the union of all entries for FNX (L/R), CNG (L/R), IFO (L/R) and SLF (L/R) cover (30.7, 25.8), (40.9, 37.2), (42.4, 46.1), (50.6, 53.3) respectively, while CST (L/R), ILF (L/R), UNC (L/R) and Fminor and Fmajor cover (23.6, 25.4), (33.4, 33.6), (14.3, 17.4), 44.3 and 34.1. Note that individual submissions appear qualitatively reasonable (Fig 2 Column 2 & 4).

**Figure 2:**
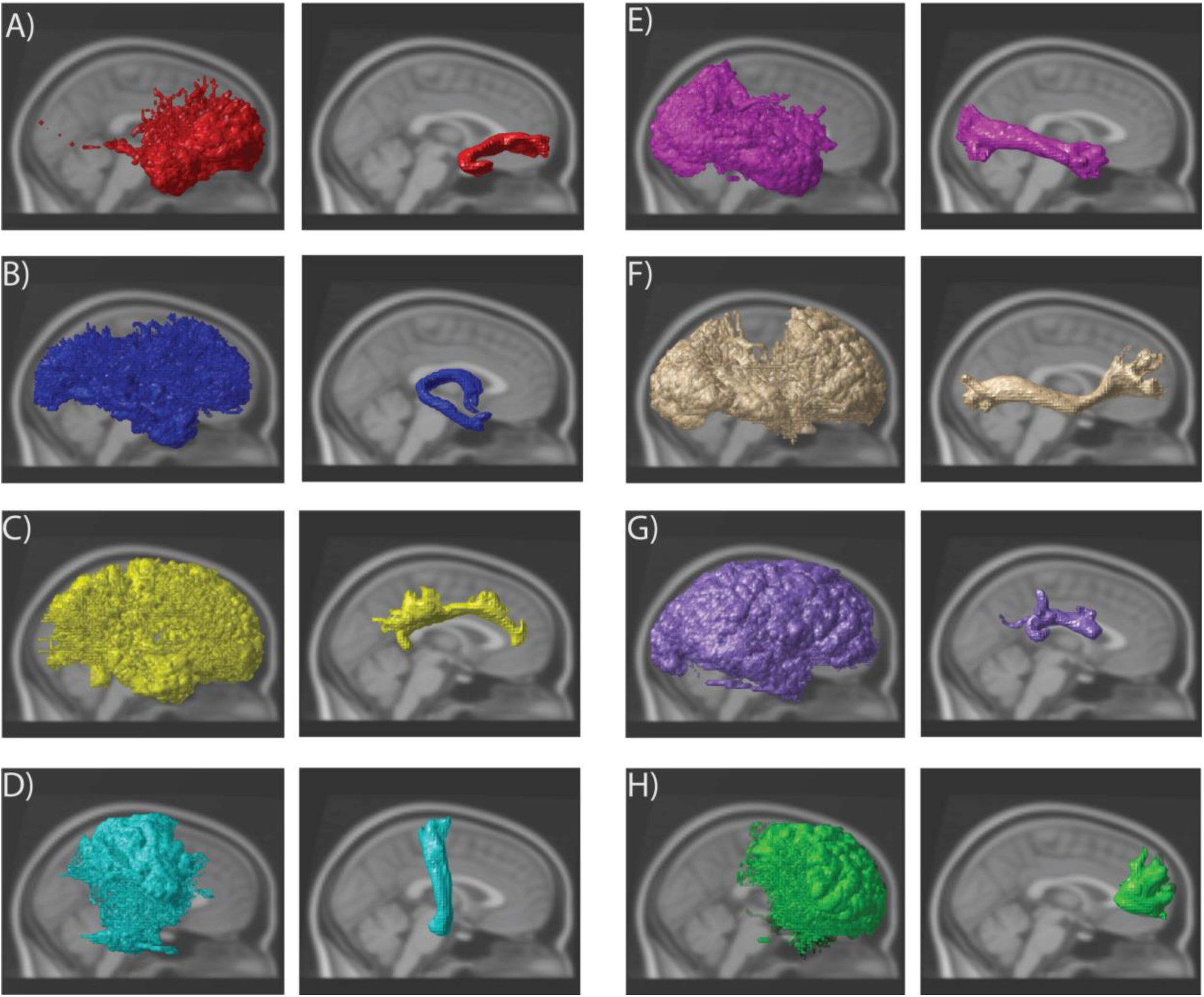
Left: An overlay of all the 46 submissions from all sessions that were acquired using both scanners per tract Right: An overlay of a single submission using all sessions that were acquired using both scanners per tract A) Uncinate left B) Fornix left C) Cingulum eft D) Corticospinal tract left E) Inferior Longitudinal Fasciculus left F) Inferior Fronto-Occipital Fasciculus left G) Superior Longitudinal Fasciculus left H) Fminor.

**Fig 3:**
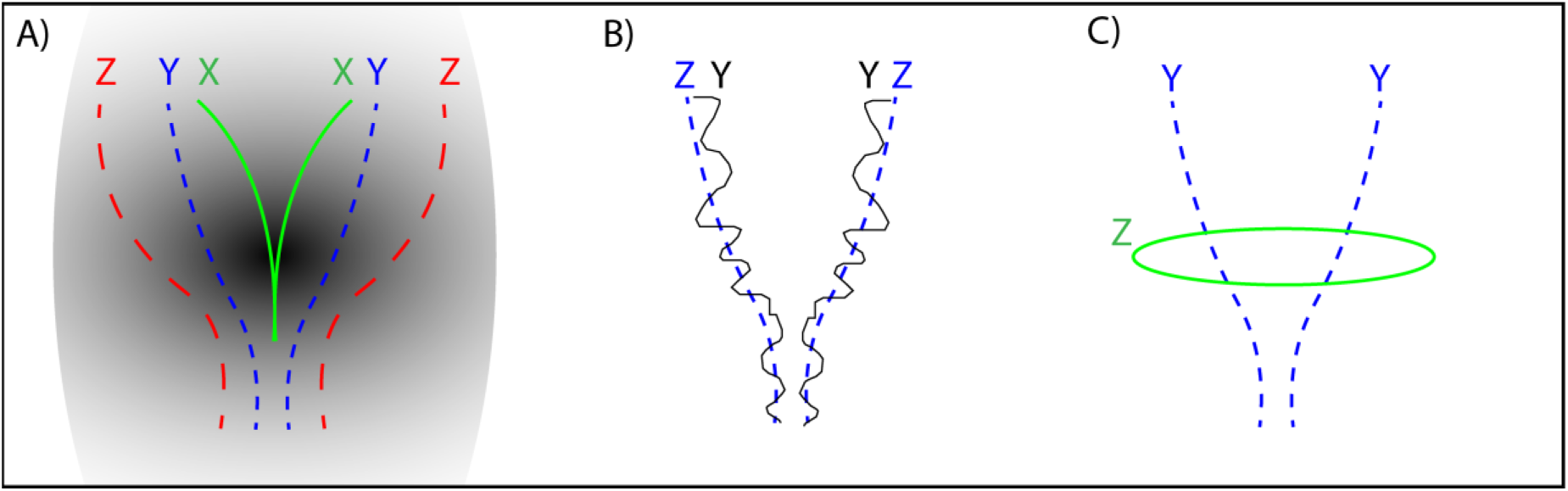
A) Where the shape X is impeccably contained in Y and Y is contained in Z. The resulting containment CI(Y, X) = 1, CI(Z, X) = 1 and CI(Z, Y) = 1. B) Shape Y is a noisy representation of shape Z where CI(Y, Z) = 0.84. C) Shape Z is different from shape Y in a different orientation and the CI(Z,Y) = 0.17

The number of algorithmic submission’s team wise are Team 1: 14, Team 2: 1, Team 3: 2, Team 4: 12, Team 5: 1, Team 6: 6, Team 7: 1, Team 8: 6 and Team 9: 3. It can be observed that the ICC range for the set of algorithms on a per team basis does not show a lot of variance. The ICC range of algorithms per team are Team 1 (0.61 – 0.77), Team 4 (0.52 – 0.58), Team 6 (0.77 – 0.85), Team 8 (0.81 – 0.89), Team 9 (0.27 – 0.69), Team 3 (0.64, 0.73), Team 2 (0.85), Team 7 (0.88) and Team 5 (0.97). The teams that submitted more than 3 algorithms show an average difference of 0.1 in terms of ICC.

Violin plots (depict the probability density of the data) of ICC and Dice for intra-session reproducibility, inter-session, and inter-scanner measures of reproducibility are presented in Figures 4, 5 and 6, respectively. Since the observations are highly similar in the afore-mentioned figures we only present a detailed comment on Figure 4 which holds true for Figure 5 and 6 as well. This figure helps in identifying the low, moderate and high reproducibility tracts. The intra-session distributions (Figure 4B) across entries for UNC (L/R) and FNX (L/R) are bi-modal with a median of the lower mode less than 0.4 ICC. The CST (L) has a smaller fraction of the entries with ICC less than 0.4, while the remainder of the entries have only a few outlier entries less than 0.4 The inter-session (Fig. 5) and inter-scanner (Fig. 6) distributions were similar, with a slight increase in outlier entries for IFO (L/R). The patterns in the dice were similar when using a quality threshold of less than 0.4 dice.

**Figure 4:**
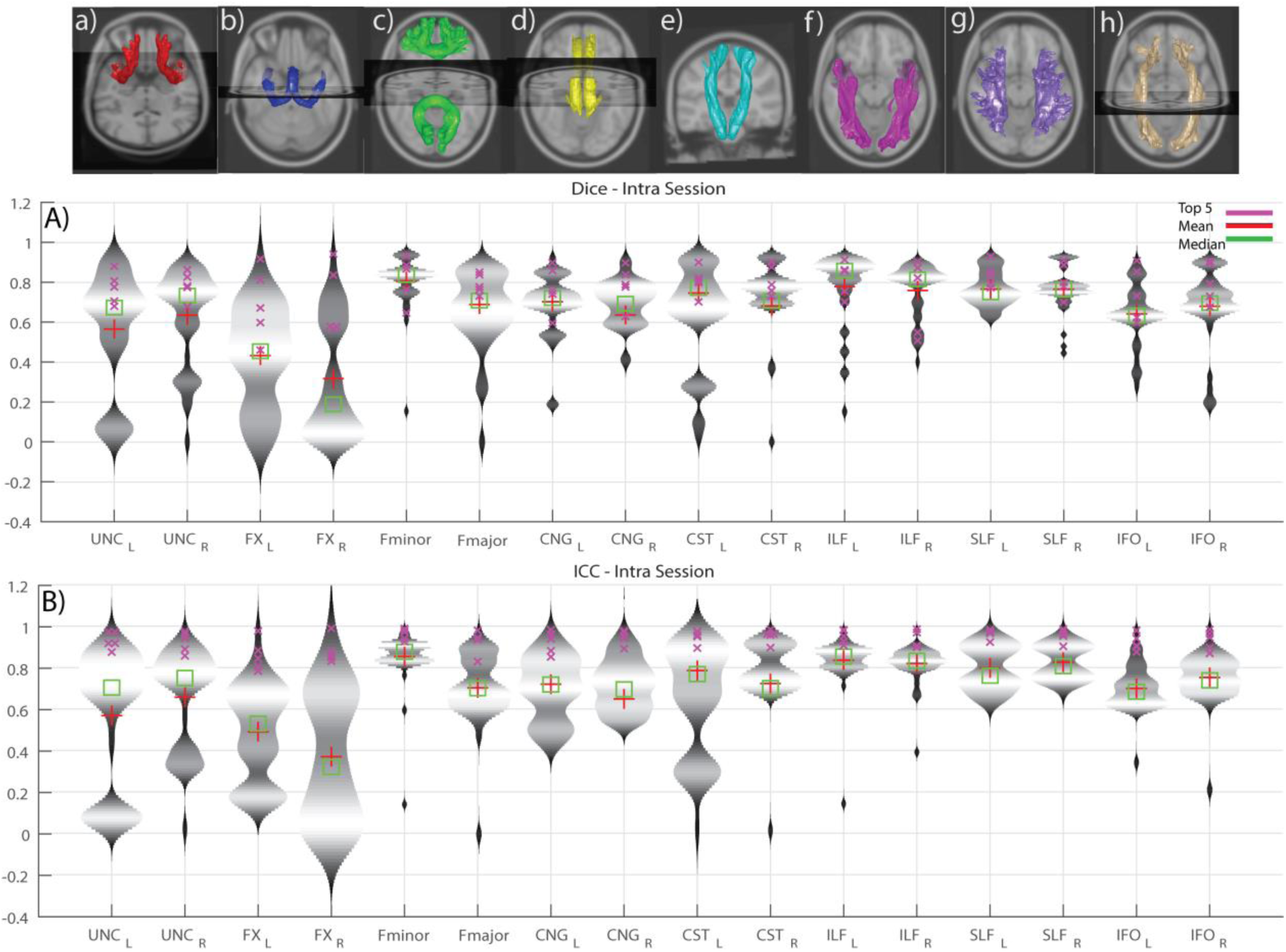
Violin plots of intra-session submissions across both the scanners per tract. A) Dice similarity coefficients B) Intra-class correlation coefficients. The top row depicts the median of the top five intra session submissions. The tracts are in the following order (L/R): a) Uncinate b) Fornix c) Fminor & Fmajor d) Cingulum e) Corticospinal tract f) Inferior longitudinal fasciculus g) Superior longitudinal fasciculus h) Inferior fronto-occipital tract

**Figure 5:**
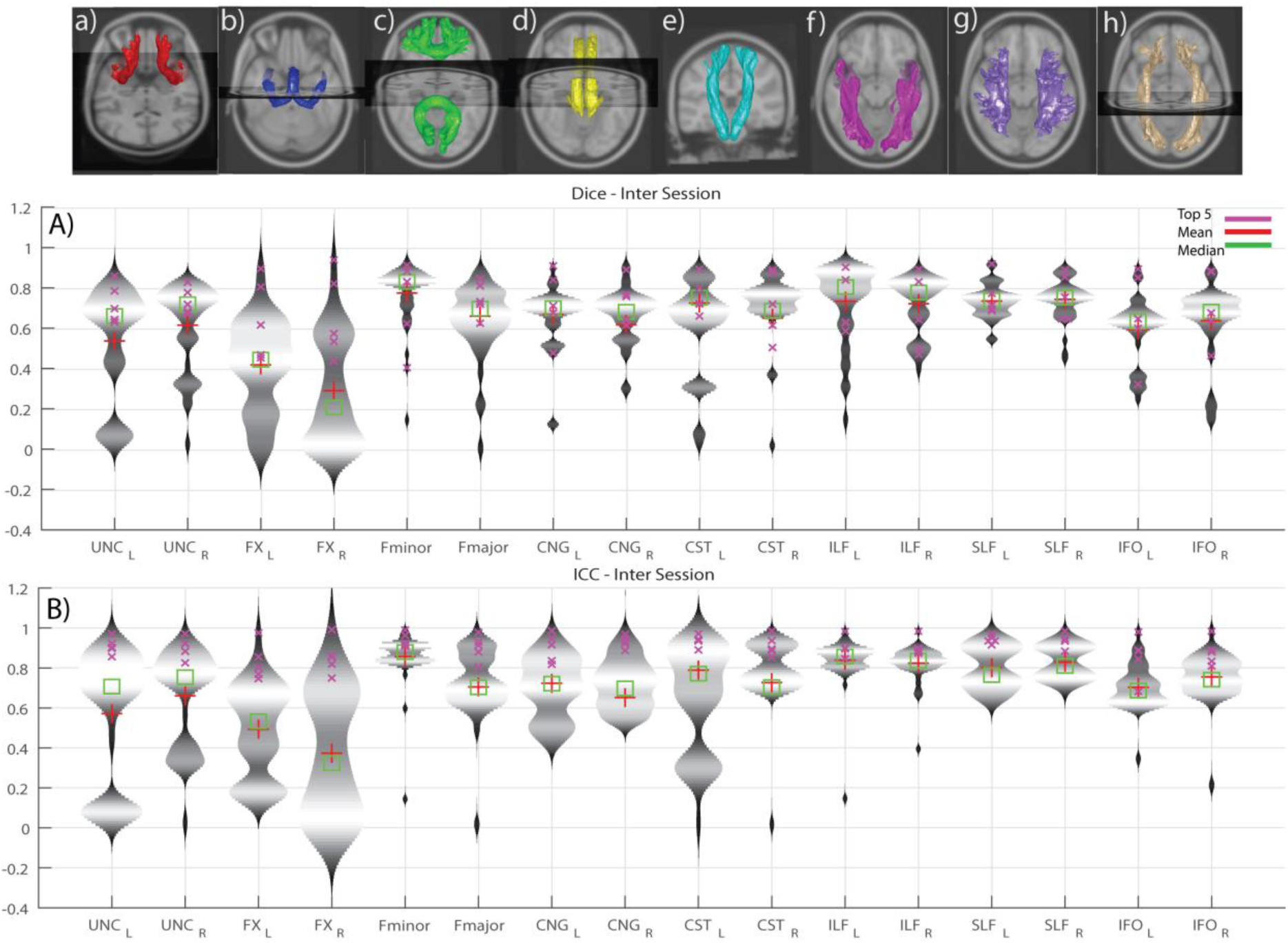
Violin plots of inter-session submissions across both the scanners per tract. A) Dice similarity coefficients B) Intra-class correlation coefficients. The top row depicts the median of the top five inter session submissions. The tracts are in the following order (L/R): a) Uncinate b) Fornix c) Fminor & Fmajor d) Cingulum e) Corticospinal tract f) Inferior longitudinal fasciculus g) Superior longitudinal fasciculus h) Inferior fronto-occipital tract

**Figure 6:**
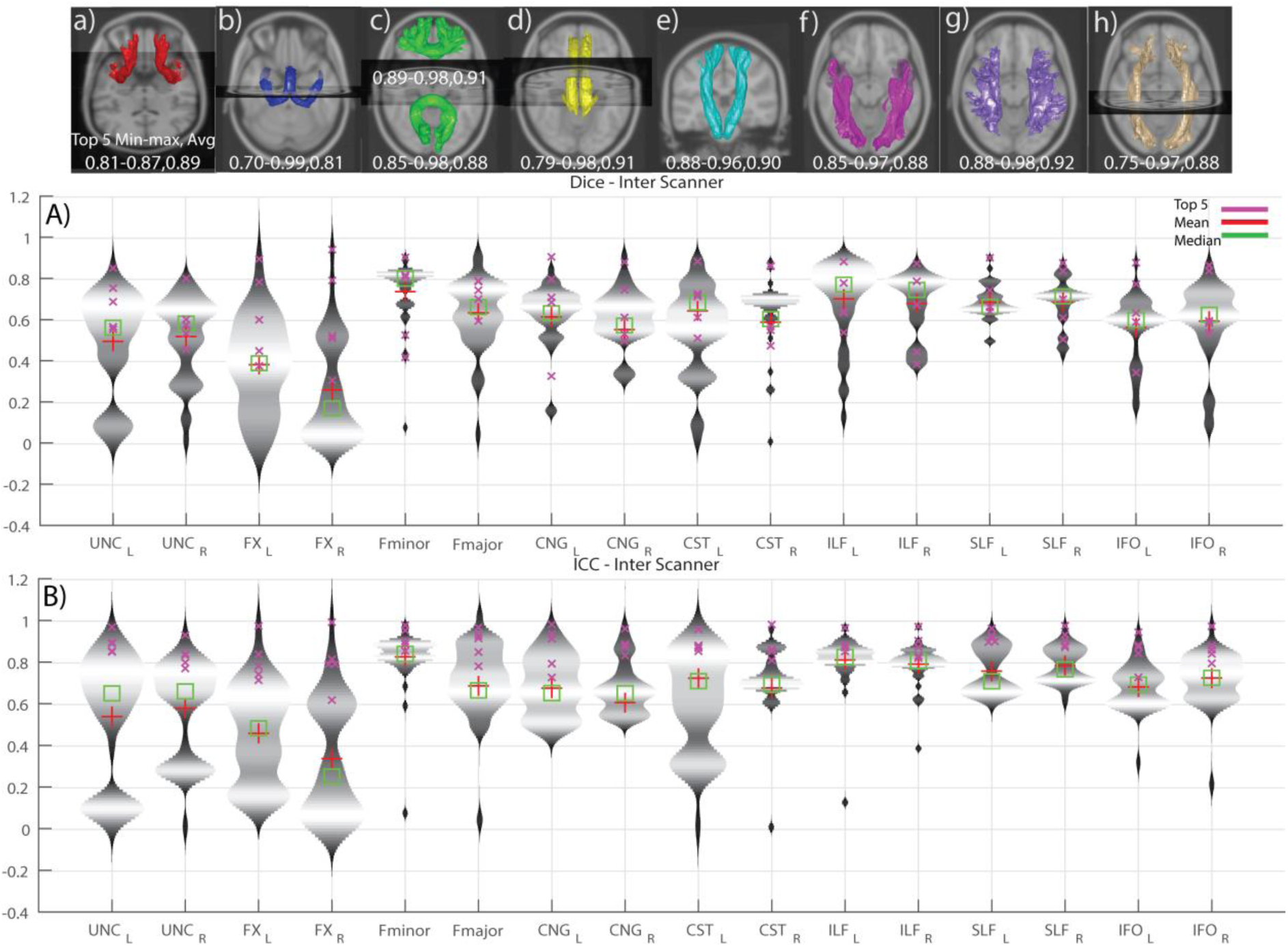
Violin plots of inter-scanner submissions across both the scanners per tract. A) Dice similarity coefficients B) Intra-class correlation coefficients. The top row depicts the median of the top five inter scanner submissions. The tracts are in the following order (L/R): a) Uncinate b) Fornix c) Fminor & Fmajor d) Cingulum e) Corticospinal tract f) Inferior longitudinal fasciculus g) Superior longitudinal fasciculus h) Inferior fronto-occipital tract

We define cutoffs for high, moderate, and low reproducibility on the inter-scanner reproducibility. High reproducibility was defined as a median ICC greater than 0.6 and less than 5% of entries less than 0.4 ICC. Moderate reproducibility was defined as median ICC greater than 0.4 and less than 25% of entries less than 0.4 ICC. Low reproducibility was defined as a median ICC less than 0.4 or more than 25% of entries less than 0.4 ICC. Hence, the high reproducibility tracts were Fminor, CST (/R), ILF (L/R), SLF (L/R) and IFO (L/R). The moderate reproducibility tracts were CST (L), Fmajor, CNG (L/R). The low reproducibility tracts were UNC (L/R) and FNX (L/R). This above is observed when looking at all submissions however when observing the top 5 submissions we see higher reproducibility.

When the analysis is restricted to only the top five submissions, we see a different picture that suggests substantively reproducible methods. The inter-scanner reproducibility among the top 5 entries in ICC (min-max, average) are shown in Fig 6.

Figure 7 illustrates the top five entries for the tracts with the lowest inter-scanner reproducibility alongside the volumetric median (median per voxel from five submissions) of the top five entries. Qualitatively, the volumetric profiles of the UNC (L/R) and FNX (L/R) are very different across the top five entries. The first submission has small “core” tracts labeled, while the second, third and fifth found much larger spatial extents and the fourth was mid-way between.

**Figure 7:**
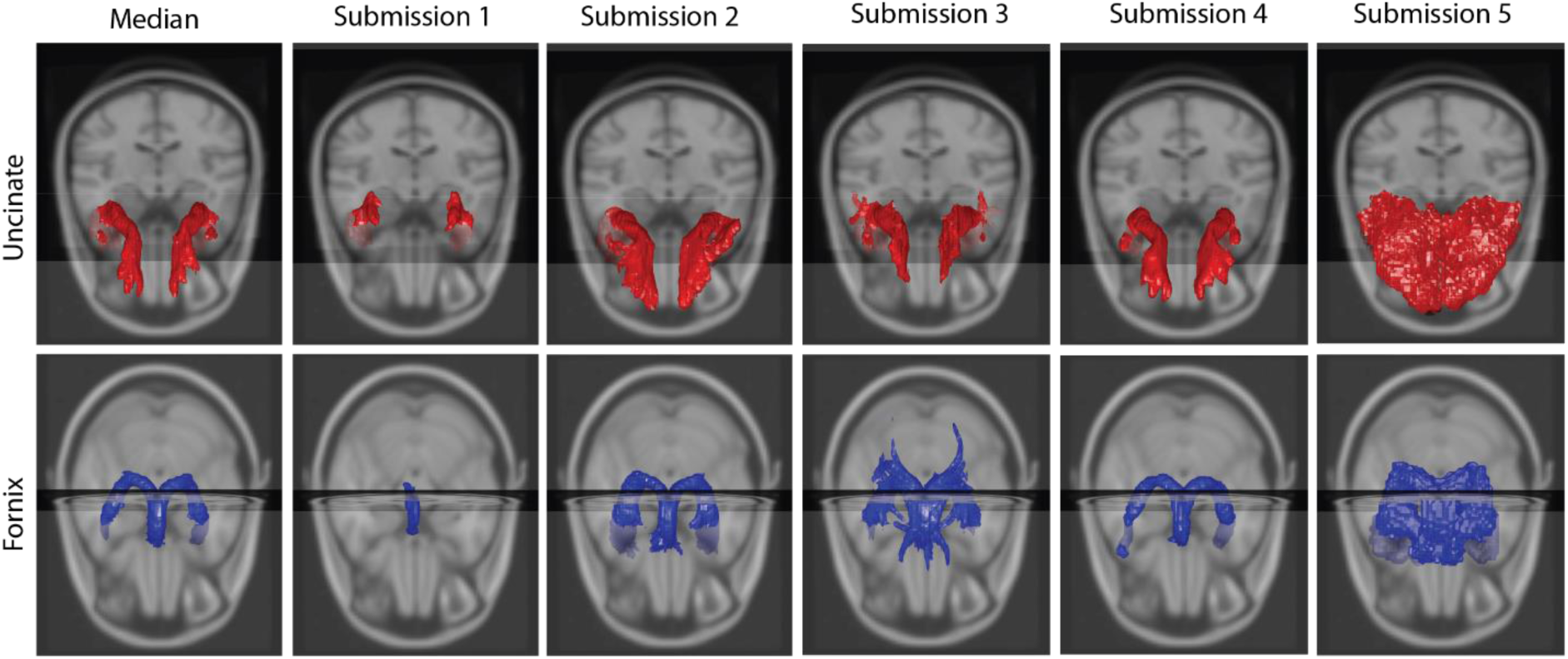
First row shows the median of Uncinate (L/R) and the top five submissions. The second row shows the median and submissions of Fornix (L/R).

## 4. DISCUSSION

The most reproducible tracts were Fminor, CST (\R), ILF (L\R), SLF (L\R), IFO (L\R), while the moderately reproducible tracts were Fmajor, CNG (L\R) and CST (\L). Lowest reproducibility tracts are UNC (L\R), FNX (L\R). These tracts have a well-spread/broad probability distribution. Note that the reproducibility of these tracts was maintained across imaging sessions and change of scanner. It is evident that all the algorithms entered are not consistently identifying the same fiber structures given the extreme variance observed in Figure 2. While most of the individual submissions show a reasonable detection of the tracts if observed from a ROI point of view (Fig 2), the difference between tract volumes between methods is quite high.

The reproducibility (ICC) of the entered algorithms varied from 0.27 to 0.97 (Fig 8A), but most of the algorithms performed with a reproducibility of 0.6 or higher. Similar levels of reproducibility were observed for methods that used selective shells or additional data from the Human Connectome Project. Note it would be inappropriate to assume independence and there are a few methods per categorical assignment, so statistical analysis across method types was not performed. Qualitatively, CSD was the most popular approach as the pre-processing fiber reconstruction method (Fig 8B). Tensor and compartment models perform well, but trailed slightly behind CSD when comparing maximum values that have been achieved using these methods. The modified version of CSD with the addition of Deep Learning U-net also performed well.

**Figure 8:**
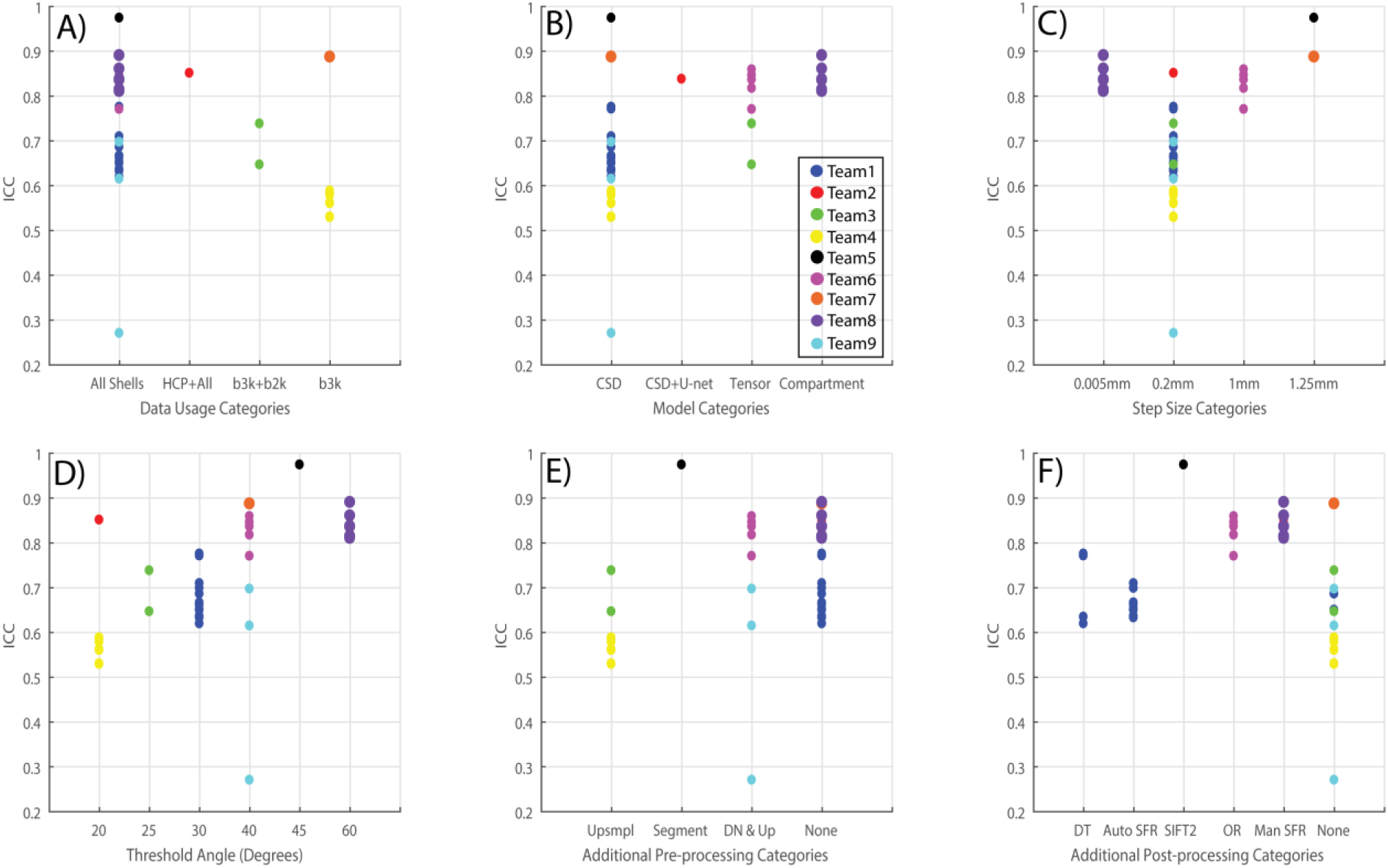
A) Quantifies the number of algorithms that used a specific part of the dataset or added more from other sources. B) Quantifies the usage of HARDI/Tensor methods by different tractography algorithms as a pre-step. C & D) Quantifies the step size and threshold angle parameter for tractography algorithms. E & F) Quantify the number of additional pre-processing and post-processing techniques applied for the tractography algorithms.

The choices of analysis parameters appears to have affected method performance. A comparison of different step sizes that have been used shows that the most heavily used category was 0.2mm (Fig 8C). However, methods using all other step size choices (e.g., 0.005, 1 and 1.25mm) performed better in terms of ICC. A variety of threshold angles have been used lying in the range of 20 – 60 degrees (Fig 8D). The variation is hard to comment upon as this suggests that a threshold angle is specific to the type of tractography algorithm. High reproducibility has been achieved at lower threshold angles such as 20 degrees and at higher angles as well such as 45 or 60 degrees. Additional pre-processing before implementing fiber reconstruction methods shows improvement for ICC only when additional segmentation was performed (Fig 8E). A comparison of de-noising coupled with up-sampling and no additional pre-processing shows higher reproducibility when no additional steps are performed. While most of the algorithms did not use additional post-processing steps (Fig 8F), the few algorithms that used the methods of outlier rejection, spurious fiber removal and SIFT2 show improvement in reproducibility. In brief, it might be inferred that additional pre-processing and post-processing techniques are helpful in increasing the reproducibility of tractography algorithms, though a systematic test of this would be necessary to draw accurate conclusions.

While it would be expected that an algorithm with empty or inaccurate bundles could achieve an extremely high ICC which would be representative of ‘null’ learning. Hence, we conducted consistency analysis using the containment index as to which bundles are contained inside which ones. The inaccurate ones will lie on the outside or show up as outliers which can be observed in Fig 9.

**Figure 9.**
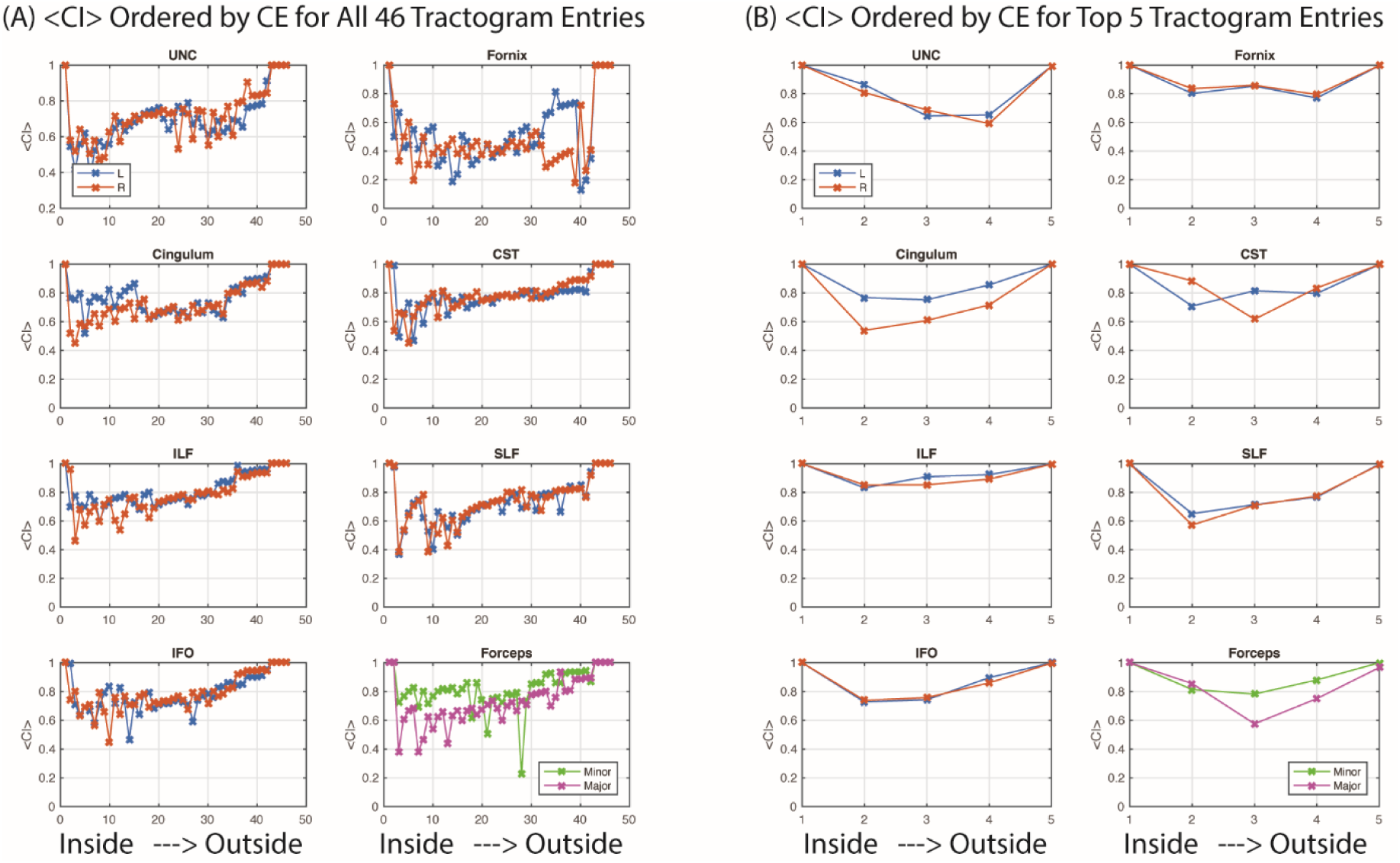
Ordering entries to minimize containment energy (CE) shows that containment index is generally lower for the volumetrically smaller tractograms (toward “inside” on each subplot) and increases for the larger tractograms (toward “outside” on each subplot). Variations in containment explained the least amount of entry variability for the UNC and Fornix, while the other tracts were more consistent. The containment between all methods (A) were more variable and lower than the containment for the top five methods (B).

As seen in Fig 9, <CI> is moderate and variable (∼0.4-0.6) for the first approximately 20 entries (after ordering) and then steadily increases for the CST, Cingulum, Forceps, ILF, IFO, and SLF. Hence, for smaller tractograms, approximately 50% of the variance is explained by nesting, but there are substantial contributions from other factors. For the larger tractograms (∼20-46 ordered entries), the differences appear largely driven by increasing volume of the tracts. UNC and Fornix are a bit more variable between ordered methods, which indicates associations within methods and suggests disagreements across major categories of entries. Finally, the Fornix is highly variables across methods (∼<0.4 <CI>), which point towards inconsistency of tract definition between approaches. When looking across all pairs of tracts, the overall rank correlation of the method ordering was low (mean=0.25) with a high variance (standard deviation=0.27, range=-0.28 to 1.0). Therefore, the relative volumetric differences between tracts were not consistent for methods across white matter tracts. Examining nestings of the top five tracts showed that Submission 5 (not shown) was always the largest, while Submission 1 and Submission 4 were determined to be the most inner methods half of the time. Submission 2 was the second largest for 12/16 tracts, while Submission 3 was the second largest for the others. This is consistent with a visualization interpretation of Figure 7. The <CI> was ∼1 for the fifth method, so a highly reproducible tract was feasible that encompassed the choices of the other top entries. The top 5 entries had high <CI> (>0.7) for the Fornix, IFO, ILF, but the remaining tracts were showed low CI for at least one method. Therefore, while at least one of the top methods differed from the others in a substantial manner, this could not be explained by volumetric differences of the tracts.

## 5. CONCLUSION

The most reproducible tracts considering all submitted algorithm outcomes are Fminor, CST (\R), ILF (L\R), SLF (L\R), IFO (L\R). The moderately reproducible ones are Fmajor, CNG (L\R) and CST (\L). Tracts with low reproducibility are UNC (L\R) and FNX (L\R). The most reproducible algorithms are 5A, 8D, 7A, 6E and 6F (Table 1) as per criteria of ICC. The mentioned algorithms are not an example of a consistent null learning as they all lie with in a nested containment with the largest covered volume.

In conclusion, the 2017 ISMRM TraCED Challenge created a publicly available multi-scanner, multi-scan in vivo reproducibility dataset and engaged nine groups with 46 algorithm entries. The TraCED Challenge dataset is freely available at www.synapse.org. Consistent with previous studies, reproducibility of tractograms was found to vary by anatomical tract. When viewed across all entries, reproducibility was concerning (ICC <0.5); however, the cluster of top performing methods resulting in reassuringly high results (ICC > 0.85). Variation in performance were seen across processing parameters, but the challenge design did not provide sufficient number of samples to identify uniformly preferred design choices. The key novel finding of this challenge is that variations in tractography methods can be largely attributed to larger/smaller volumetric difference tradeoffs for the larger tracts, especially among methods that are tuned towards volumetrically larger tractograms. Yet, the different methods clearly result in fundamentally different tract structures at the more conservative specificity choices (i.e., volumetrically smaller tractograms). The containment index, containment energy, and containment index framework provides a consistent approach to evaluate the nesting structure tractograms, and the freely available data and results from this challenge can be used to quantify new tractography approaches.

## ACKNOWLEDGEMENTS

This work was supported by R01EB017230 (Landman). This work was conducted in part using the resources of the Advanced Computing Center for Research and Education at Vanderbilt University, Nashville, TN. This project was supported in part by the National Center for Research Resources, Grant UL1 RR024975-01, and is now at the National Center for Advancing Translational Sciences, Grant 2 UL1 TR000445-06. The content is solely the responsibility of the authors and does not necessarily represent the official views of the NIH. This work is supported in part by China Scholarship Council Scholarship. This work was supported in part by the National Natural Science Foundation of China (Grant No. 61379020).

